# Lipoxin A_4_/FPR2 signaling mitigates ferroptosis of alveolar epithelial cells via NRF2-dependent pathway during lung ischemia-reperfusion injury

**DOI:** 10.1101/2024.04.22.590127

**Authors:** Denny Joseph Manual Kollareth, Victoria Leroy, Zhenxiao Tu, Makena Jade Woolet-Stockton, Manasi Kamat, Timothy J. Garrett, Carl Atkinson, Guoshuai Cai, Gilbert R. Upchurch, Ashish K. Sharma

## Abstract

**BACKGROUND:** Post-lung transplantation (LTx) injury can involve sterile inflammation due to ischemia-reperfusion injury (IRI). We investigated the cell-specific role of ferroptosis (excessive iron-mediated cell death) in mediating lung IRI and determined if specialized pro-resolving mediators such as Lipoxin A4 (LxA_4_) can protect against ferroptosis in lung IRI.

**METHODS:** Single-cell RNA sequencing of lung tissue from post-LTx patients was analyzed. Lung IRI was evaluated in C57BL/6 (WT), formyl peptide receptor 2 knockout (*Fpr2^−/−^*) and nuclear factor erythroid 2-related factor 2 knockout (*Nrf2^−/−^*) mice using a hilar-ligation model with or without LxA_4_ administration. Furthermore, the protective efficacy of LxA_4_ was evaluated employing a murine orthotopic LTx model and *in vitro* studies using alveolar type II epithelial (ATII) cells.

**RESULTS:** Differential expression of ferroptosis-related genes was observed in post-LTx patient samples compared to healthy controls. A significant increase in the levels of oxidized lipids and reduction in the levels of intact lipids were observed in mice subjected to IRI compared to shams. Furthermore, pharmacological inhibition of ferroptosis with liproxstatin-1 mitigated lung IRI and lung dysfunction. Importantly, LxA_4_ treatment attenuated pulmonary dysfunction, ferroptosis and inflammation in WT mice subjected to lung IRI, but not in *Fpr2^−/−^* or *Nrf2^−/−^*mice, after IRI. In the murine LTx model, LxA_4_ treatment increased PaO_2_ levels and attenuated lung IRI. Mechanistically, LxA_4_-mediated protection involves increase in NRF2 activation and glutathione concentration as well as decrease in MDA levels in ATII cells.

**CONCLUSIONS:** LxA_4_/FPR2 signaling on ATII cells mitigates ferroptosis via NRF2 activation and protects against lung IRI.

## INTRODUCTION

Lung transplantation (LTx) is an established option for patients with end stage lung diseases after other therapeutic measures have been exhausted (1). Survival after LTx still lags far behind that observed after other solid organ transplantations, as evidenced by a median survival rate that currently stands at 5.8 years (2). Post LTx ischemia–reperfusion injury (IRI), which occurs during the process of donor lung preservation and transplantation, contributes to the development of primary graft dysfunction (PGD) and chronic lung allograft dysfunction (CLAD) (3). During IRI, the endogenous resolution of inflammation mechanisms become compromised, leading to prolonged and dysregulated inflammation (4). Hallmarks and sequalae of lung IRI include increased acute innate immunity responses and oxidative stress resulting in inflammation, vascular permeability, pulmonary edema, alveolar damage and ultimately lung dysfunction (5).

Resolution of acute inflammation is a biochemically active process, regulated by endogenous mediators called specialized pro-resolving mediators (SPMs), that act in concert to switch off the inflammatory response and return the tissue to homeostasis (6). SPMs are produced by cells of the innate immune system, which are formed via the stereoselective conversion of polyunsaturated fatty acids that include arachidonic acid, eicosapentaenoic acid and docosahexaenoic acid. They are grouped into four families, lipoxins, resolvins, protectins, and maresins (7). Lipoxin A_4_ (LxA_4_), a member of the lipoxin family, is produced from the metabolism of arachidonic acid, and has been reported to exert a variety of activities in multiple tissues, including anti-inflammatory effects, regulation of neutrophil infiltration, pro-resolving signaling, macrophage polarization, and uptake of apoptotic polymorphonuclear neutrophils (8). LxA_4_ functions as a “braking signal” in the inflammatory response and switches the inflammatory response to the resolution phase (9). The activities of LxA_4_ are regulated by the G protein-coupled formyl peptide receptor 2 (FPR2) and have been shown to protect against ferroptosis in neuronal pathology(10). However, the role of ferroptosis in lung IRI and the efficacy of LxA_4_ to attenuate ferroptosis in lung IRI remains to be delineated.

Ferroptosis is an iron-dependent, non-apoptotic form of cell death, which mediates its effects in part by promoting the accumulation of lethal lipid ROS (11). Biochemically, the mechanism underlying ferroptosis is mainly related to glutathione (GSH) depletion, inactivation of glutathione peroxidase 4 (GPX4), iron overload and lipid peroxidation (12). Ferroptosis has been implicated in pathological cell death associated with all kinds of diseases, including degenerative diseases, carcinogenesis, and IRI, and attenuation of ferroptosis has been shown to reduce lipopolysaccharide induced lung injury (13, 14).

Our previous study determined that SPMs such as LxA_4_ are significantly and sequentially increased in human post-LTx bronchoalveolar lavage (BAL) samples (4). In this study, we investigated the cell-specific role of ferroptosis in lung IRI and elucidated the effectiveness of LxA_4_ to mitigate lung IRI by protecting against ferroptosis-mediated inflammation. Our findings show that LxA_4_ attenuated ferroptosis and inflammation in lung IRI mediated through FPR2 and NRF2-dependent signaling pathways. Collectively, our findings suggest that administration of exogenous LxA_4_ would serve as a promising therapeutic option for lung IRI.

## MATERIALS AND METHODS

Detailed Methods are described in the Supplementary Materials.

### Lung IRI model

An *in vivo* hilar-ligation model of lung IRI was used in 8 to 12-week-old C57BL/6 (WT), formyl peptide receptor 2 knockout (*Fpr2*^−/−^) and nuclear factor erythroid 2-related factor 2 (*Nrf2^−/−^*) mice, as previously described (15).

### Deceased Cardiac Donor (DCD) and Orthotopic Lung Transplantation

LTx were performed utilizing cuff techniques using Balb/c donor and C57BL/6 recipient mice, as previously described (15)

## RESULTS

### Differential expression of ferroptosis-related genes in ATII cells of CLAD patients

We performed single-cell RNA sequencing analysis to investigate if ferroptosis plays a contributing role in post-LTx patients with chronic allograft graft dysfunction (CLAD). Our analysis utilized a previously reported study of scRNA sequencing analysis from 4 CLAD patients and 3 donor controls (donor tissue; DT) (16). We identified 6,046 ATII cells and 6,099 ATI cells and focused on these cells in downstream analysis (**Figure 1A**).

**Figure 1.**
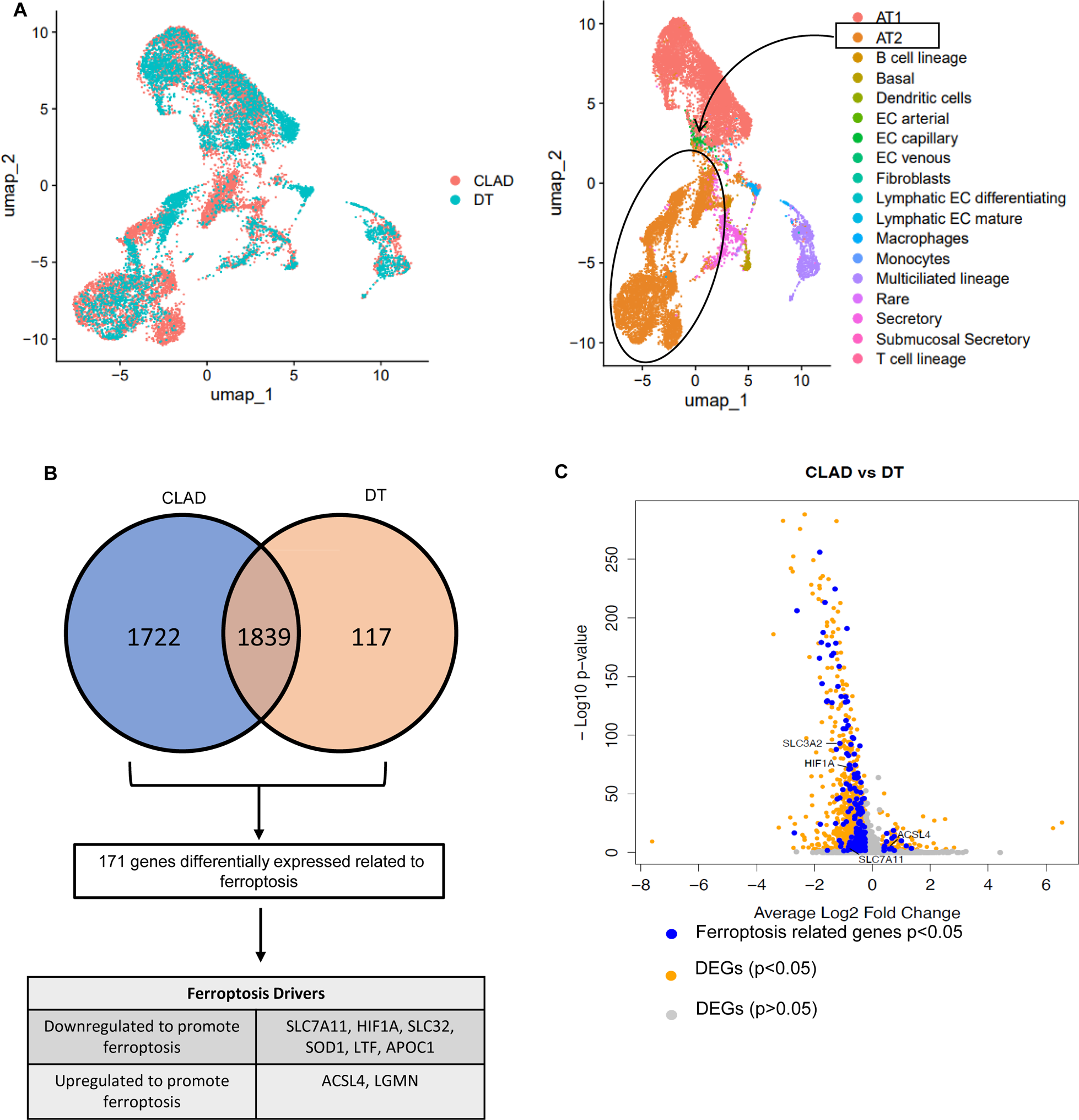
Single cell RNA sequencing analysis reveal differential regulation of ferroptosis related genes in ATII cells of CLAD patients. (**A**) UMAP visualization illustrating X cell clusters within lung tissue samples from both CLAD patients and donor control (**B**) Venn diagram depicting the identification of differentially expressed genes (DEGs) associated with ferroptosis. The diagram highlights downregulated genes in CLAD (blue) and donor tissue (DT; yellow). (**C**) Volcano plot illustrating the differentially expressed genes (DEGs) within ATII cells. Ferroptosis related genes (FRGs) with a significant differential expression (p<0.05) are shown by blue dots while other significant DEGs (p<0.05) are represented by orange dots. Grey dots represent genes that are not statistically significant (p>0.05).

In depth analysis of 1839 differentially expressed genes (DEGs) revealed that, out of the top 50 downregulated DEGs in CLAD, 12 were related to ferroptosis (LGALS1, TNFAIP3, EMP1, TIMP2, SNHG6, MYH9, GRN, SH3BGRL3, CTSZ, CDKN1A, IRF1, PLAUR) (**Figure 1B**). Moreover, of the top 50 upregulated DEGs, 6 were related to ferroptosis (SFTA1P, SLC39A8, SPCS2, PDIA6, PDIA4, LGMN) (surveyed from genes outlined in **Table S1-S2**). Additionally, we observed alterations in key ferroptosis related genes in human CLAD. Specifically, there was significant downregulation of SLC7A11, HIF1A, SLC3A2 and upregulation of ACSL4 in CLAD when compared to DT (**Figure 1C**). The expression of SLC7A11, SLC3A2 and HIF1A are known to inhibit ferroptosis and downregulation of these genes indicate increased ferroptosis in human CLAD (17, 18). Furthermore, ACSL4 is a ferroptosis inducer (19) and overexpression of ACSL4 indicates ongoing ferroptosis in patients with CLAD compared to DT controls. We also observed similar changes in regulation of ferroptosis related genes in ATI cells (**Supplementary Figure S1**). Collectively, these results suggest a pivotal association of chronic graft rejection with involvement of ferroptosis in epithelial cells of post-LTx patient cohorts.

### Ferroptosis is involved during lung IRI

Intracellular iron accumulation, increased levels of oxidized lipid products and subsequent membrane destabilization are key events in initiating ferroptosis (20). In the present study, we determined whether these key signaling molecules involved in ferroptosis are associated with lung IRI (**Figure 2A**). Using the murine hilar ligation model of lung IRI, we observed that expression of total iron, Fe^2+^ and Fe^3+^ levels were significantly higher in lung tissue of mice subjected to IRI compared to sham (**Figure 2B**). A marked increase in ferric ion staining was also observed in lung tissue after IRI compared to sham (**Supplementary Figure S2**). Moreover, lipidomic analysis of lung tissue showed that there was a significant increase in the levels of oxidized lipids in mice subjected to IRI compared to sham. Specifically, we found significantly increased levels of oxidized phosphatidylcholine (OxPC), oxidized phosphatidylethanolamine (OxPE), and oxidized triglycerides (OxTG) that correlated with increase in lung dysfunction and IRI. Furthermore, there was a significant reduction in the levels of intact lipid species such as PC, PE, plasmenyl-PC, plasmenyl-PE and sphingomyelin (SM) as well as lysophospholipids in mice lungs subjected to IRI compared to sham (**Figure 2C-D and Table S3**). Taken together, these results suggest that ferroptosis-specific lipids are involved in the pathogenesis of lung IRI.

**Figure 2.**
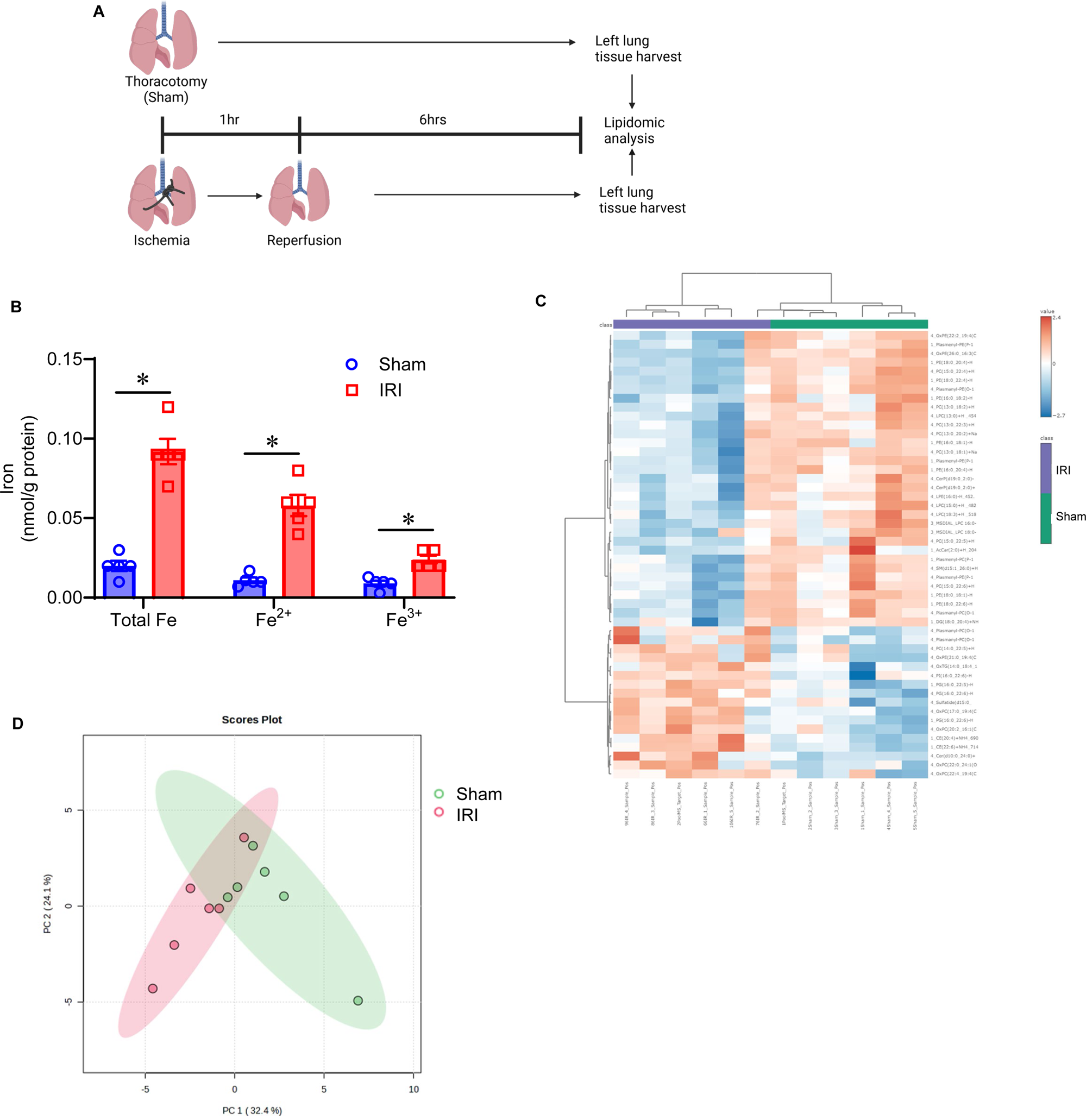
Increased ferroptosis is observed in lung tissues of mice subjected to IRI. (**A**) Schematic depicting the left lung hilar ligation model of murine IRI. (**B**) Significant increase in excessive iron levels were observed in mice subjected to IRI compared to the sham group; *p< 0.009; n=5/group. (**C**) Heatmap from untargeted lipidomic depicting increased levels of oxidized lipids and decreased levels of intact lipids in lung IRI; n=6/group; Ox, oxidized; PI, phosphatidylinositol; TG, triglycerides; PG, phosphatidylglycerol; PE, phosphatidylethanolamine; PC, phosphatidylcholine; CE, cholesteryl ester; AcCar, acyl carnitine; LPC, lysophosphatidylcholine; DG, diacylglycerols; SM, sphingomyelin; LPE, lysophosphatidylethanolamine; Cer, Ceramide. (**D**) Principal component analysis (PCA) to determine main differences in lipid profiles between sham and IRI groups.

### Pharmacological inhibition of ferroptosis alleviates lung IRI

We next assessed whether pharmacological inhibition of ferroptosis can attenuate lung dysfunction and IRI by using Liproxstatin-1, a specific and potent inhibitor of ferroptosis, in the murine IRI model. A significant increase in pulmonary dysfunction is observed after lung IRI (6hrs), as evidenced by increased airway resistance, pulmonary artery (PA) pressure and decreased pulmonary compliance, compared to sham controls (**Figure 3A-D**). Importantly, mice treated with Liproxstatin-1 offered significant protection against lung dysfunction compared to vehicle-treated mice. Furthermore, we observed significant increases in hallmarks of ferroptosis, such as depleted GSH levels and increased MDA levels, in lung tissue of mice subjected to IRI compared to sham (**Figure 3E-F**). Liproxstatin-1 treatment reduced ferroptosis as evidenced by significant increase in GSH levels and decrease in MDA levels compared to vehicle-treated mice. Additionally, pharmacological inhibition of ferroptosis significantly attenuated the expression of pro-inflammatory cytokines such as TNF-α, IL-17A, CXCL-1, RANTES, IL-6 and MIP-1α in BAL fluid of mice undergoing lung IRI compared to untreated controls, suggesting a crosstalk between ferroptosis and inflammation during lung IRI (**Figure 3G-L**). To further explore the protective role of ferroptosis inhibition on lung IRI, neutrophil infiltration in lung tissue and MPO levels in BAL fluid were quantified. A marked increase in neutrophil infiltration was observed in WT mice subjected to IRI, while Liproxstatin-1 treatment significantly reduced neutrophil infiltration in lung tissue and myeloperoxidase expression in BAL fluid compared to untreated mice (**Figure 3M-O**).

**Figure 3.**
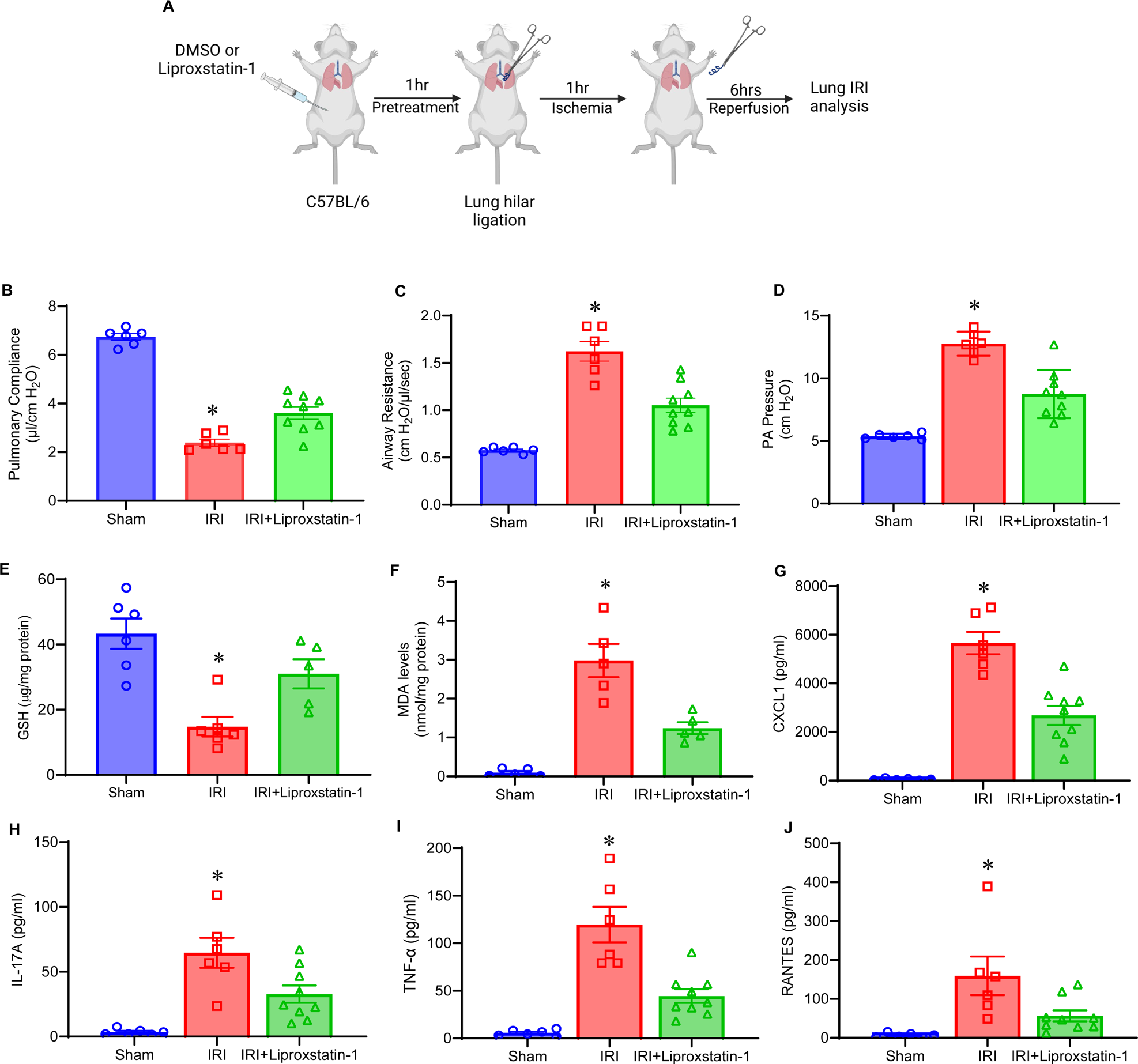
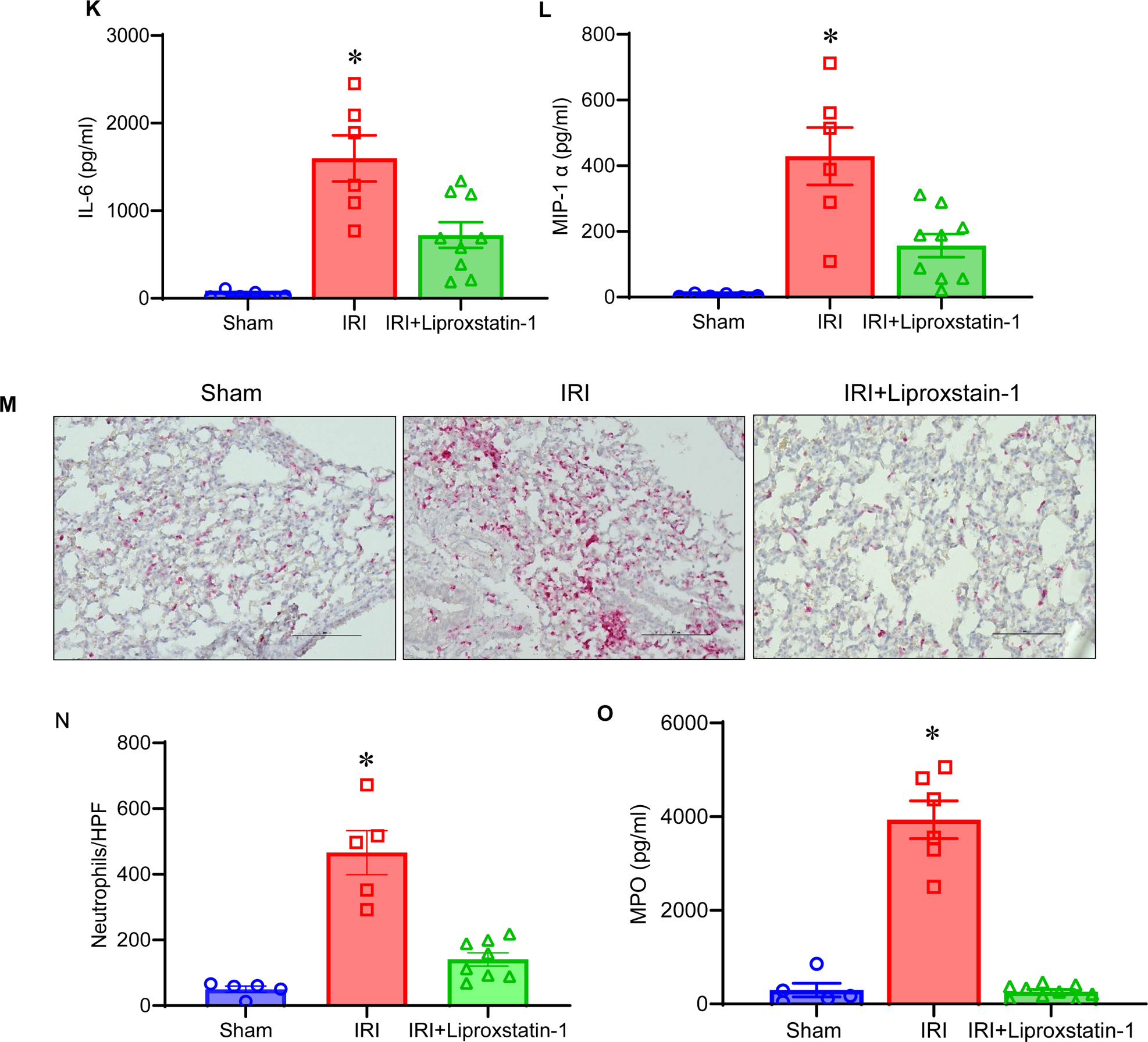
Pharmacological inhibition of ferroptosis alleviates lung IRI. (**A**) Schematic depicting the liproxstatin-1 treatment in the murine lung IRI model. (**B-D**) Pre-treatment with liproxstatin-1 offered significant protection in lung dysfunction as shown by increased pulmonary compliance, decreased airway resistance and PA pressure; *p<0.01 vs. other groups; n=6-9/group. (**E-F**) Lung IRI induced depletion of GSH and increase in MDA was mitigated by Liproxstatin-1 treatment to decrease ferroptosis in lung tissue. *p<0.01 vs. other groups; n=6-9/group. (**G-L**) Pro-inflammatory cytokine and chemokine levels were significantly attenuated in BAL fluid after Liproxstain-1 treatment in mice subjected to IRI compared to untreated mice. *p<0.03 vs other groups; n=6-9/group. (**M**) Representative images of neutrophil staining from mice lung sections after sham or IRI treated with or without Liproxstain-1; Scale bars indicate 100 µm. (**N-O**) Neutrophil infiltration and MPO levels were significantly reduced in mice treated with Liproxstatin-1 compared to untreated mice. *p<0.03 vs. other groups; n=5-9/group.

### Lipoxin A_4_ protects against pulmonary dysfunction and ferroptosis via FPR2 signaling

To assess the influence of specific bioactive isoforms of SPMs and their receptors on regulation of ferroptosis during lung IRI, exogenous LxA_4_ was administered to WT and *Fpr2^−/−^* mice and assessed for lung function and IRI (**Figure 4A**). WT mice treated with LxA_4_ showed significant improvement in lung function as evidenced by increase in pulmonary compliance, decrease in airway resistance and PA pressure (**Figure 4B-D**). However, LxA_4_ treatment attenuated lung dysfunction in WT mice, but not in *Fpr2^−/−^* mice after lung IRI, as observed by decrease in pulmonary compliance, as well as increases in airway resistance and PA pressure (**Figure 4B-D**). Moreover, administration of LxA_4_ reduced ferroptosis in WT mice, but not in *Fpr2^−/−^* mice, as shown by an increase in GSH content and decrease in MDA levels in WT mice compared to *Fpr2^−/−^* mice (**Figure 4E-F**). Additionally, LxA_4_ treated WT mice, but not LxA4-treated *Fpr2^−/−^* mice, demonstrated significant mitigation of pro-inflammatory cytokines and chemokines compared to respective controls (**Figure 4G-L**). Similarly, neutrophil infiltration and MPO levels following IRI was mitigated in LxA_4_ treated WT mice, but not *Fpr2^−/−^*mice, compared to respective controls (**Figure 4M-O**).

**Figure 4.**
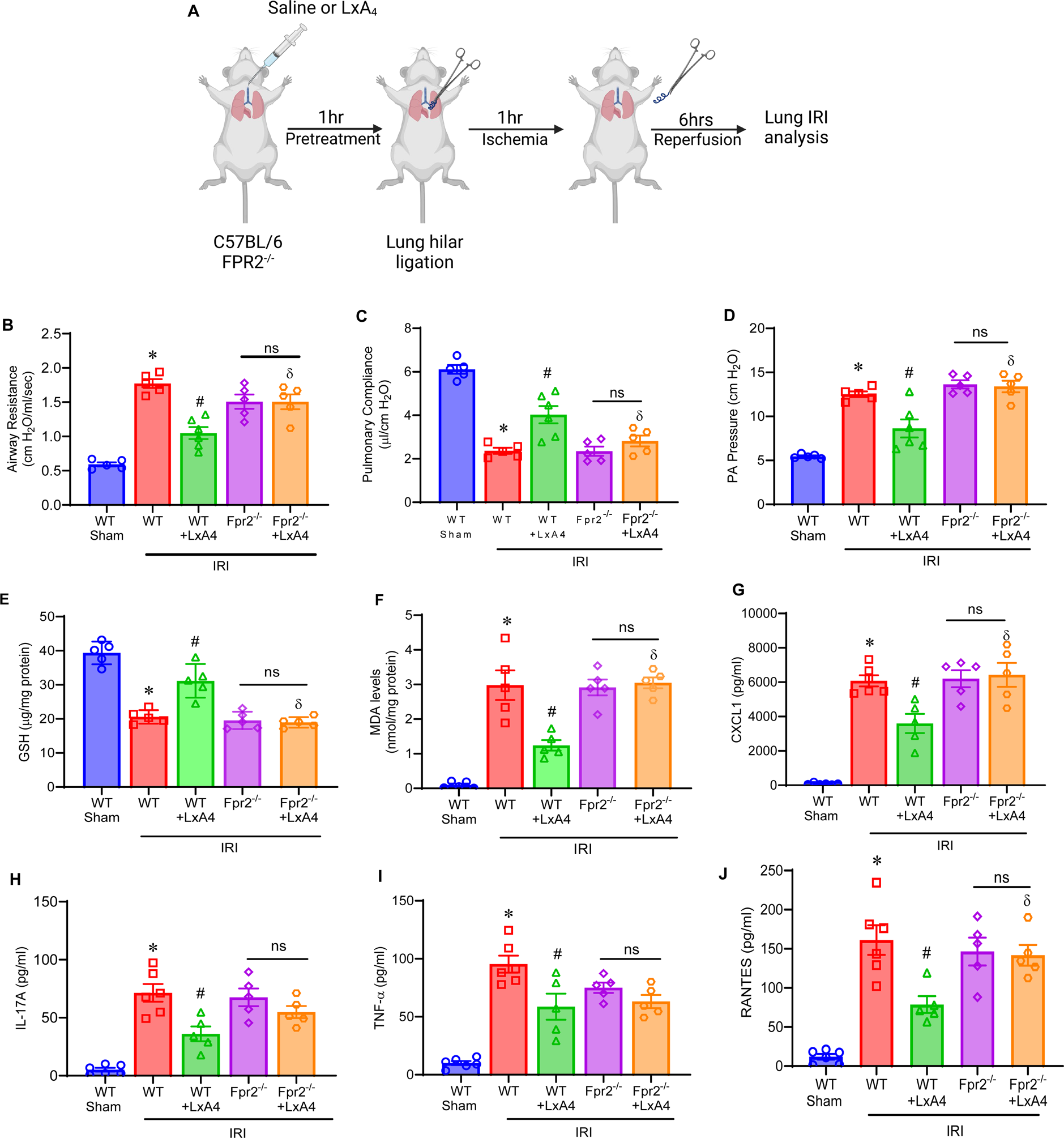
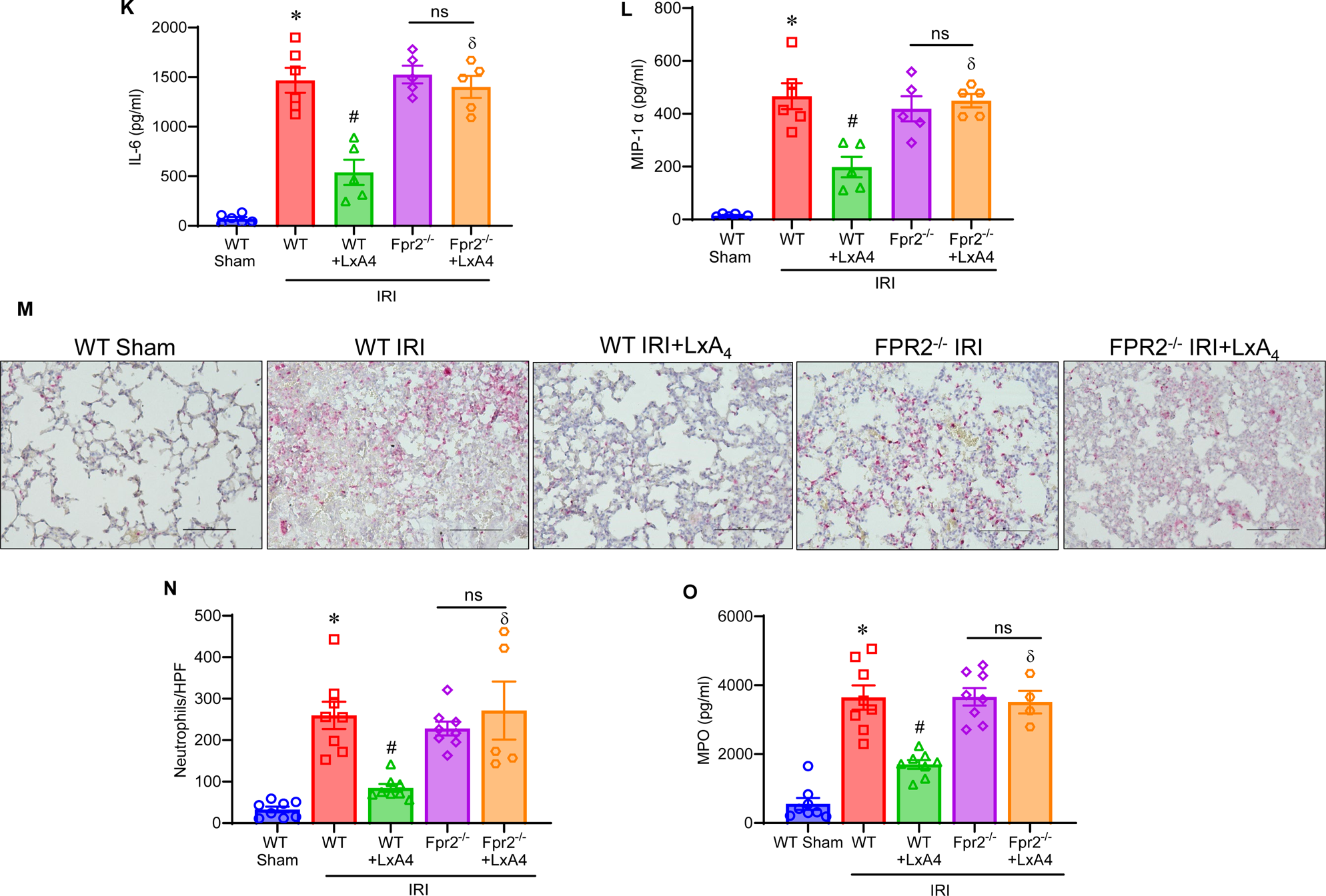
LxA_4_ mediated attenuation of pulmonary dysfunction, IRI and ferroptosis is FPR2 dependent. (**A**) Schematic representation of LxA_4_ treatment in mouse lung IRI model using WT and *Fpr2^−/−^* mice. (**B-D**) LxA_4_ treatment reduced pulmonary dysfunction following IRI in mice compared to untreated mice; *p<0.0001 vs. sham; #p<0.002 vs. WT IRI; ^δ^p<0.01 vs. WT IRI+LxA4; ns, non-significant; n=5-6/group. (**E-F**) LxA_4_ reduced ferroptosis in lungs after IRI in WT mice, but not in *Fpr2^−/−^* mice, as shown by increased GSH and decreased MDA levels. *p<0.0001 vs sham; #p<0.0004 vs WT IRI; ^δ^p<0.01 vs. WT IRI+LxA4; ns, non-significant; n=5/group. (**G-L**) LxA_4_ administration significantly reduced inflammatory cytokine and chemokine levels in BAL following IRI in WT mice, but not in *Fpr2^−/−^* mice, compared to untreated mice. *p<0.0001 vs sham; #p<0.006 vs WT IRI; ^δ^p<0.03 vs. WT IRI+LxA_4_; ns, non-significant; n=5-6/group. (**M**) Representative images of neutrophil staining in mice. Scale bars indicate 100 µm. (**N-O**) LxA_4_ treatment significantly reduced neutrophil infiltration and MPO levels in BAL fluid in mice subjected to IRI in WT mice, but not in *Fpr2^−/−^* mice, compared to untreated mice. *p<0.001 vs. sham; #p<0.001 vs. IRI; ^δ^p<0.03 vs. WT IRI+LxA_4_; ns, not significant; n=5-6/group.

### Lipoxin A_4_ reduces lung IRI by activating NRF2 pathway

NRF2 is a master regulator of the antioxidant response and has been shown to regulate the activity of several ferroptosis and lipid peroxidation-related proteins (21). Therefore, we investigated whether LxA_4_/FPR2 signaling exerts its protective actions through the activation of NRF2 in lung IRI (**Figure 5A**). We found that LxA_4_ administration offered protection against IRI mediated pulmonary dysfunctions in WT mice, but not in *Nrf2^−/−^* mice, as evidenced by decreased pulmonary compliance and increased airway resistance and PA pressure (**Figure 5B-D**). Additionally, LxA_4_ decreased ferroptosis in lung tissue (**Figure 5E-F**) and inflammation in BAL fluid (**Figure 5G-L**) of WT mice, but not in *Nrf2^−/−^* mice. Similarly, we observed significant reduction in neutrophil infiltration and MPO levels in WT mice treated with LxA_4,_ but not in *Nrf2^−/−^* mice (**Figure M-O**).

**Figure 5.**
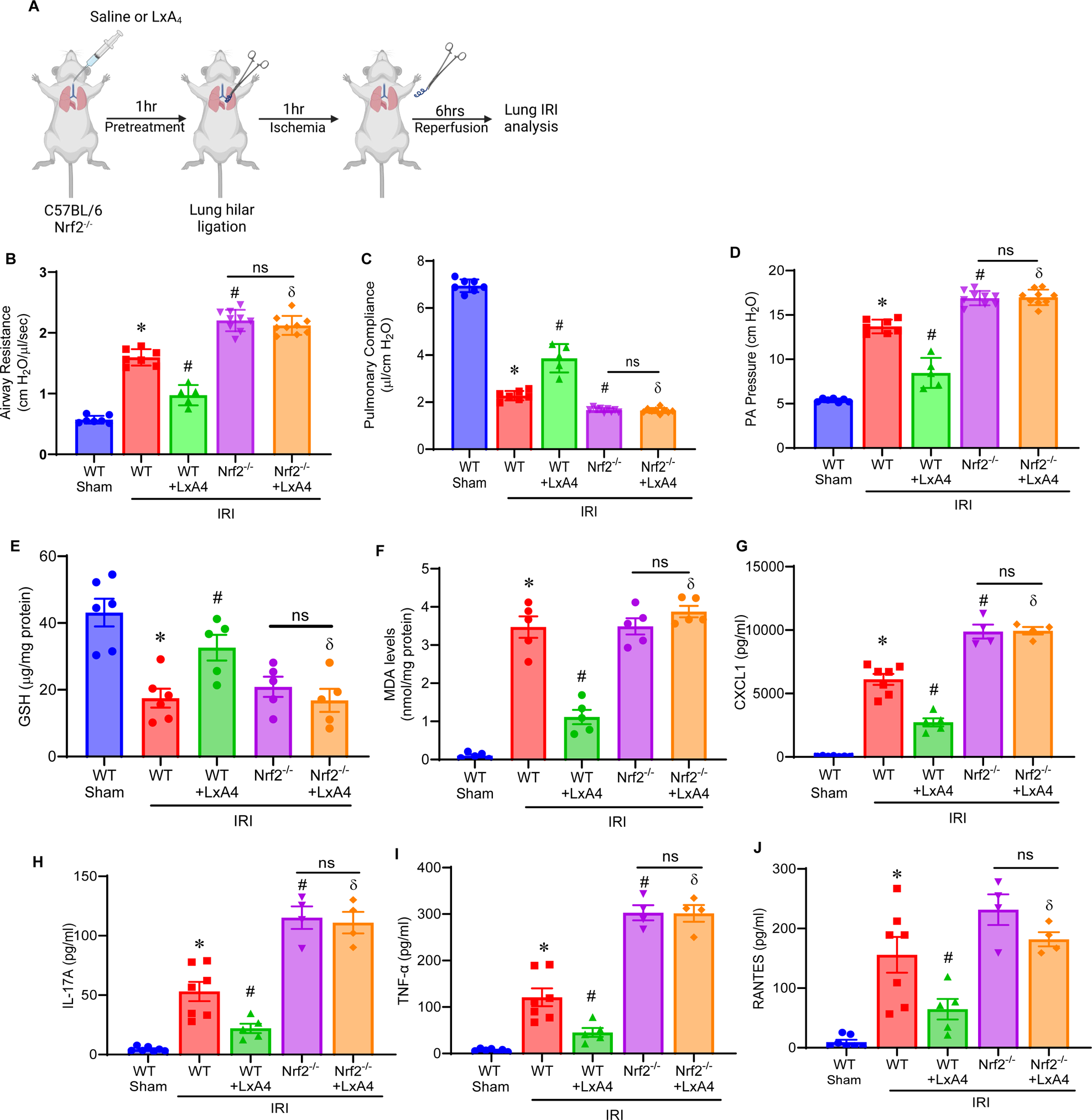
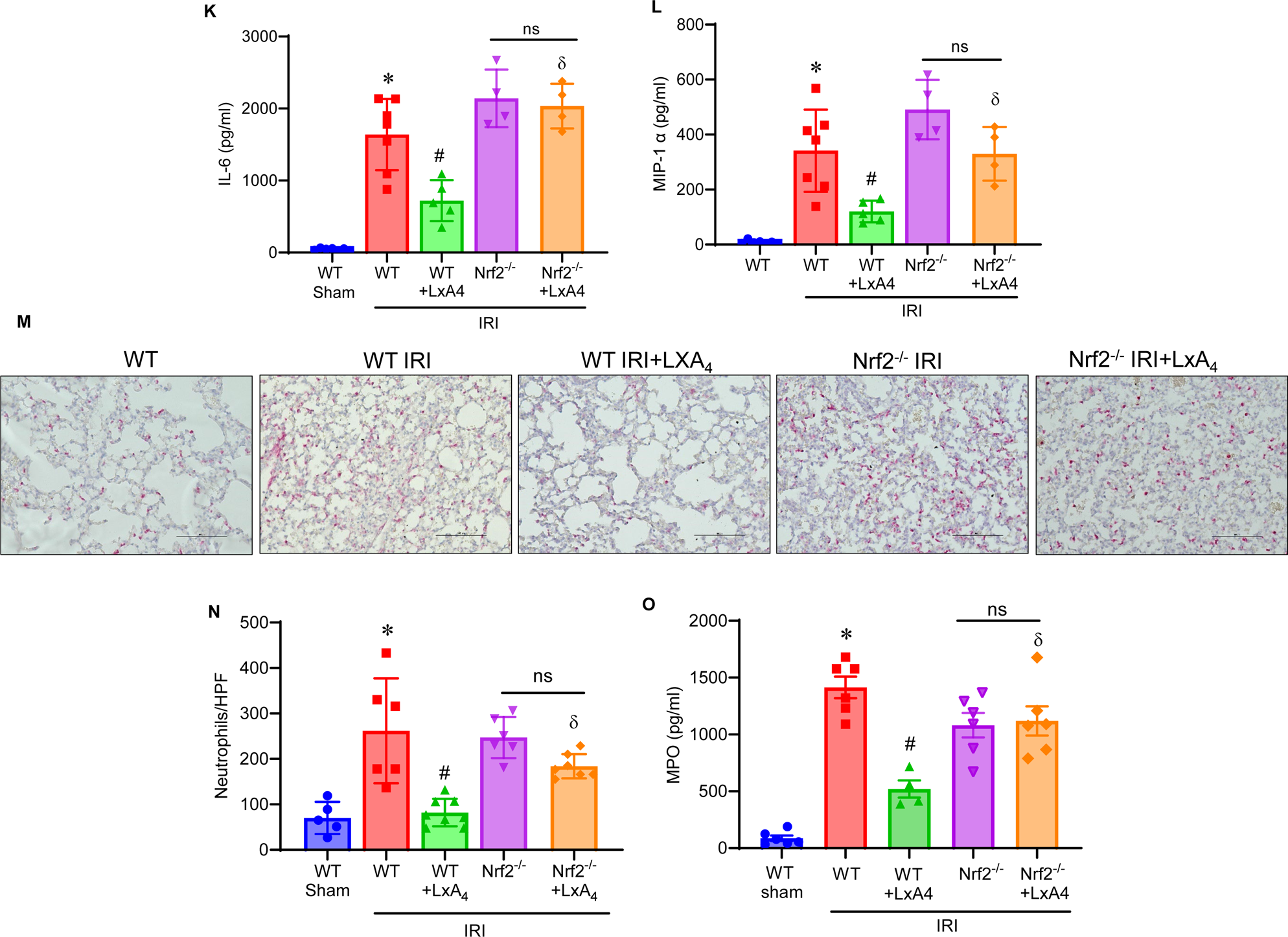
LxA_4_ reduces lung injury and ferroptosis via modulation of NRF2. **(A)** Schematic representation of mouse lung IRI model using WT and *Nrf2^−/−^* mice. (**B-D**) LxA_4_ treatment protected against IRI-induced pulmonary dysfunction as evidenced by increased pulmonary compliance, decreased airway resistance and PA pressure in WT mice, but not in *Nrf2^−/−^* mice. *p<0.0001 vs sham; #p<0.0001 vs WT IRI; ^δ^p<0.01 vs. WT IRI+LxA_4_; ns, non-significant; n=5-9/group (**E-F**) LxA_4_ treatment reduced ferroptosis in WT mice, but not in *Nrf2^−/−^* mice, compared to untreated mice after IRI. *p<0.0001 vs. sham; #p<0.002 vs. WT IRI; ^δ^p<0.01 vs. WT IRI+LxA_4_; ns, non-significant; n=5-6/group. (**G-L**) LxA_4_ treatment reduced pro-inflammatory cytokine and chemokine levels in WT mice subjected to IRI, but not in *Nrf2^−/−^* mice; *p<0.0001 vs sham; #p<0.03 vs WT IRI; ^δ^p<0.02 vs WT IRI+LxA_4_; ns, non-significant; n=4-7/group (**M**) Representative histology images of neutrophil staining in murine lung sections. (**N-O**) Neutrophil infiltration and MPO levels were significantly reduced in LxA_4_ treated WT mice, but not in *Nrf2^−/−^* mice, compared to untreated mice. n=5-6/group. *p<0.008 vs sham; #p<0.02 vs WT IRI; ^δ^p<0.01 vs. WT IRI+LxA_4_; ns, non-significant; n=4-7/group.

### Post-transplant lung injury is significantly ameliorated by LxA_4_ treatment in DCD donors

Post-LTx involves tissue injury due to characteristics of donor allografts as well as cold storage that can impact lung IRI. Therefore, to confirm our results from the hilar-ligation IRI model, we determined the impact of LxA_4_ treatment in an experimental murine orthotopic lung transplant model using donation after circulatory death (DCD) lungs. Donor mice were treated with LxA_4_ or normal saline (vehicle control) immediately upon DCD induction and recipient mice were treated immediately before LTx (**Figure 6A**). A significant increase in PaO_2_ levels in LxA_4_ treated mice was observed compared to vehicle treated controls (**Figure 6B**). The levels of pro-inflammatory cytokines and chemokines were significantly reduced in LxA_4_-treated mice compared to untreated mice (**Figure 6C-H**). Furthermore, LxA_4_ treatment demonstrated significant reduction in neutrophil infiltration and MPO levels compared to control lungs after DCD (**Figure 6I-K**). Additionally, albumin levels were significantly reduced in LxA_4_ treated mice compared to controls (**Figure 6L**). Together, these findings show the role of LxA_4_ in modulating inflammation and improving lung functions following lung IRI.

**Figure 6.**
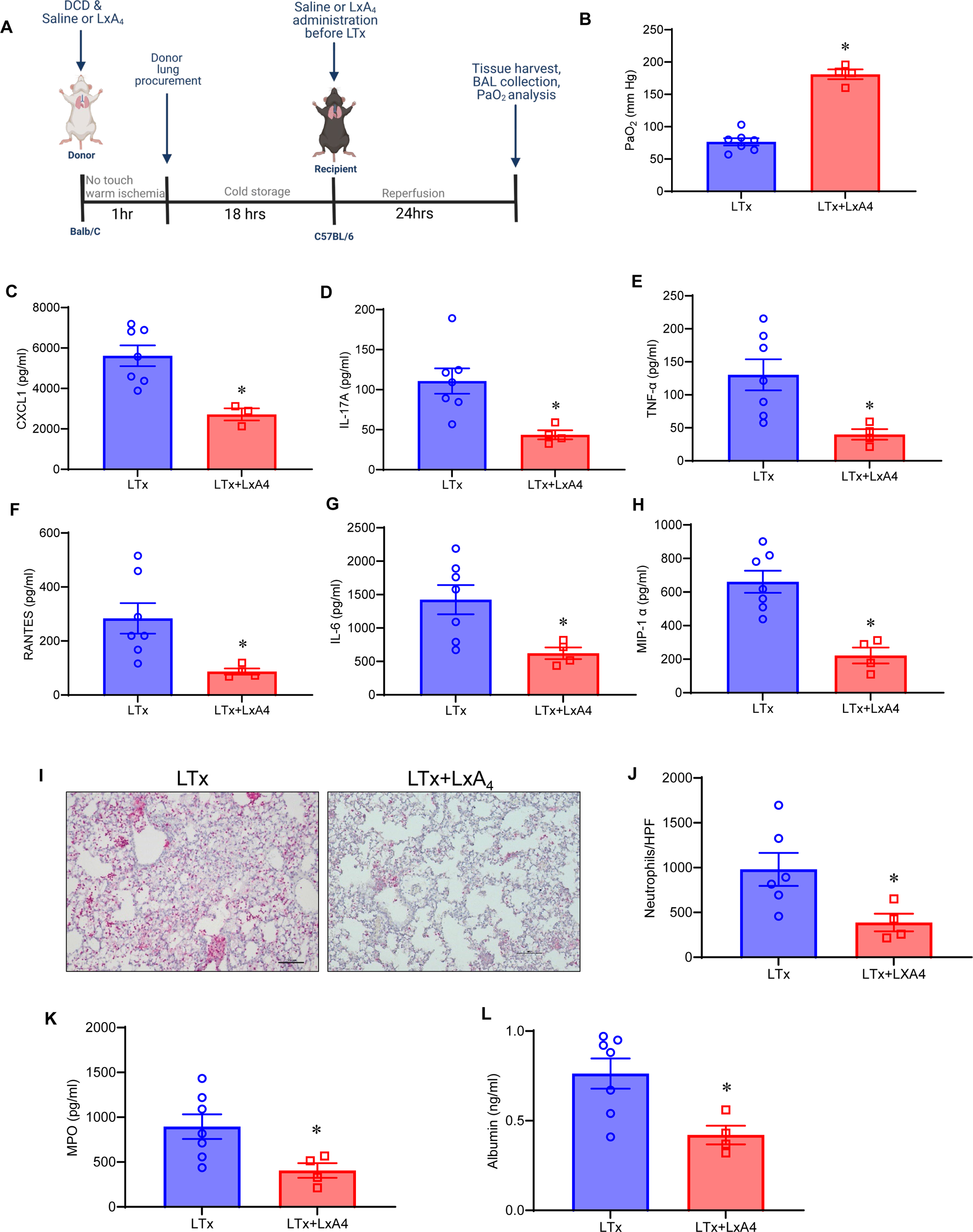
LxA_4_ reduces lung allograft IRI in recipients of DCD lungs. (**A**) Schematic representation of LxA_4_ treatment in an orthotopic murine LTx model. (**B**) LxA_4_ treatment significantly increased PaO_2_ levels in LTx mice compared to vehicle treated controls; *p<0.0001 vs LTx; n=4-7/group (**C-H**) Pro-inflammatory cytokines and chemokine expression in BAL fluid were significantly reduced in LTx mice treated with LxA_4_ compared to untreated mice. *p<0.04 vs LTx; n=3-7/group (**I**) Representative images of histological immunostaining of neutrophils of murine lung sections. (**J-K**) LxA_4_ treatment significantly abrogated neutrophil infiltration and MPO levels after post-LTx IRI compared to untreated mice. *p<0.04 vs LTx; n=5/group. (**L**) LxA_4_ treatment significantly mitigated lung edema as measured by albumin content in BAL, compared to untreated mice. *p<0.04 vs LTx; n=5/group.

### LxA_4_ reduces ferroptosis in lung epithelial cells via activation of NRF2

ATII cells are one of the critical target cells for lung IRI that are involved in neutrophil chemotaxis and their dysfunction is a vital effector process of lung injury (22). *In vitro* studies were performed using ATII epithelial cells to investigate cell specific signaling mediated by LxA_4_ in a FPR2-dependent manner (**Figure 7A**). ATII cells subjected to HR+cytomix (TNF-α and IL-17A) treatment demonstrated significantly decreased GSH, and increased MDA levels and reactive oxygen species (ROS) (**Figure 7B-E**). Importantly, LxA_4_ treatment significantly increased GSH and NRF2 activation as well as attenuated MDA and ROS levels to suppress ferroptosis (**Figure 7B-E**). A significant increase in MDA expression and decrease in GSH levels was observed in erastin (ferroptosis activator)-treated ATII cultures which was inhibited by LxA4 treatment (**Supplementary Figure S3**). Moreover, LxA_4_-mediated protection against ferroptosis of ATII cells was regulated by FPR2 receptors, as FPR2-siRNA transfected cells did not mitigate the HR+cytomix induced decrease in GSH or increase in MDA or ROS after LxA_4_ treatment compared to c-siRNA transfected cells (**Figure 7B-E**). Taken together, the pivotal role of LxA_4_ in mitigating ferroptosis in ATII cells ultimately leads to decreased CXCL1 secretion and neutrophil transmigration in a FPR2-dependent manner, thereby leading to attenuation of lung IRI (**Figure 7F**).

**Figure 7.**
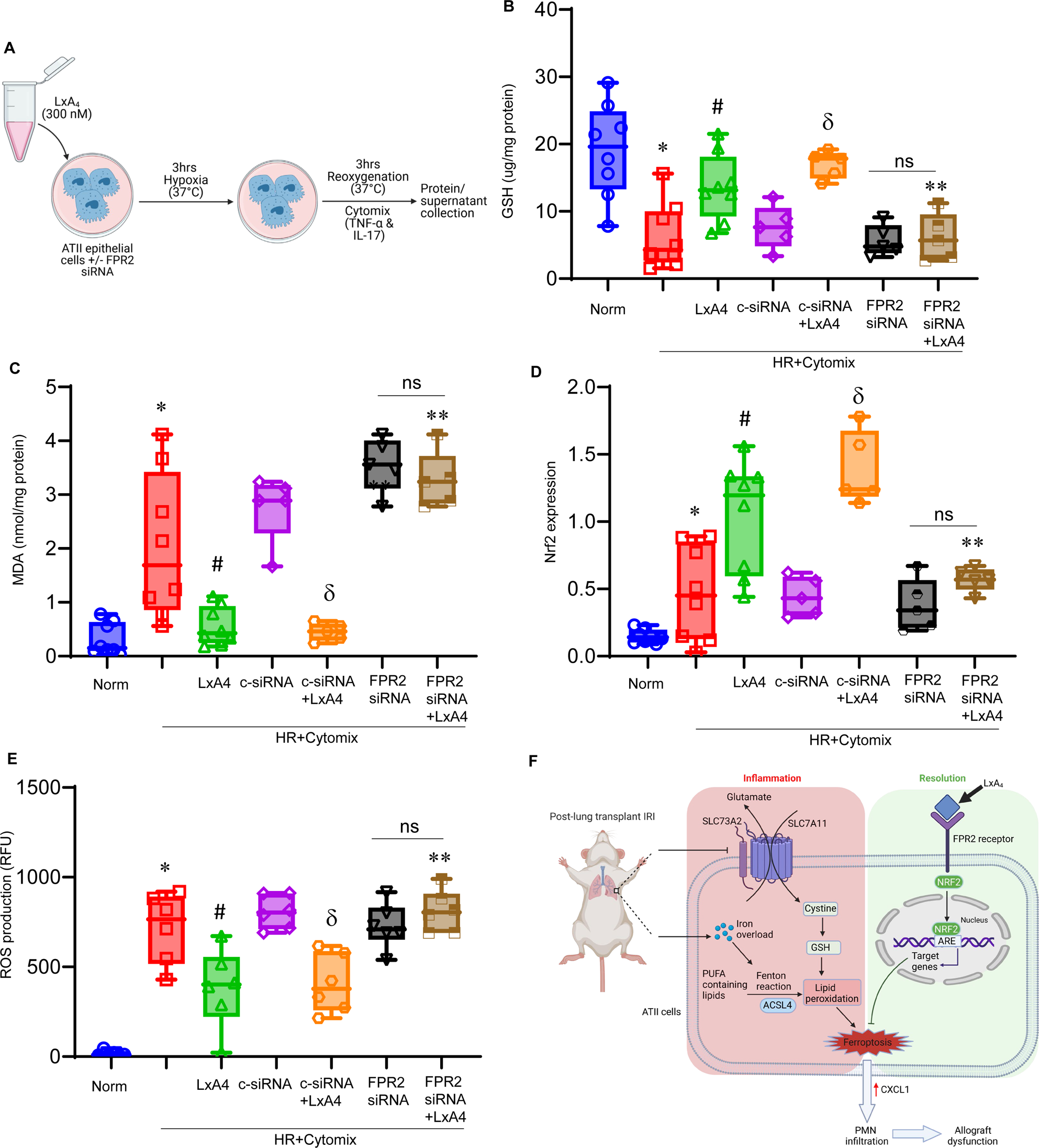
LxA_4_ treatment reduces ferroptosis in ATII cells via NRF2 activation. (**A**) Schematic showing *in vitro* analysis of LxA_4_-mediated protection in ATII cells. (**B-C**) HR+cytomix treatment induced depletion of GSH and increase in MDA levels that was decreased by LxA_4_ treatment. Moreover, HR+cytomix induced decrease in GSH concentration and increase in MDA levels which was mitigated by LxA_4_ treatment in c-siRNA treated cells but nor in FPR2 siRNA-treated cells. (**D**) LxA_4_ treatment of ATII cells increased the nuclear expression of NRF2 after HR+cytomix treatment compared to untreated cells alone in an FPR2-dependent manner. (**E**) ROS production was assessed in ATII epithelial cells by quantification of fluorescence intensity in cells treated with dichlorofluorescein (DCF) dye. HR+cytomix-induced increase in ROS production was attenuated by LxA_4_ treatment in a FPR2-dependent manner. *p<0.01 vs. Normoxia (norm); ^#^p<0.01 vs. HR+cytomix; ^δ^p<0.02 vs. c-siRNA+HR+cytomix; **p<0.03 vs. c-siRNA+LxA_4_+HR+cytomix; ns, not significant. n=5-8/group. (**F**) Schematic depicting the protective mechanism mediated by LxA_4_/FPR2 signaling in the lung microenvironment following IRI. LxA_4_ treatment mitigates ferroptosis in ATII cells via NRF2 activation, that culminates in decreased CXCL1-mediated chemotaxis and attenuation of neutrophil transmigration, leading to mitigation of pulmonary inflammation and protection against IRI and allograft dysfunction.

## DISCUSSION

Recent studies have suggested a pivotal role of SPMs in limiting the inflammatory response, promoting the resolution of inflammation, and maintaining homeostasis after tissue injury. Specific SPMs have been demonstrated to have remarkable anti-inflammatory properties leading to inflammation-resolution by regulating the activation of immune cells, and mediating the crosstalk with parenchymal cells to limit the injury cascade (23). Previously, we have shown that there is a significant increase in LxA_4_ levels in human BAL from post LTx patients suggesting that LxA_4_ plays an important role in endogenous mechanisms that facilitate resolution of inflammation (4). The results reported in the current study describe a previously uncharacterized role of LxA_4_ in attenuating ferroptosis after lung IRI via mechanistic signaling on ATII cells. Our study found that ferroptosis is involved in lung IRI in the early stages of reperfusion and attenuation of ferroptosis reduces inflammation and improves IRI induced pulmonary dysfunctions. Importantly, exogenous administration of LxA_4_ mitigates ferroptosis, lung dysfunction and neutrophil infiltration via an FPR2-dependent mechanism to attenuate post-LTx IRI. Mechanistically, LxA_4_/FPR2 signaling increases NRF2 activation of ATII cells to downregulate ferroptosis and suppression of CXCL1 release that leads to impediment of neutrophil transmigration from the vasculature to the alveolar space, thereby attenuating post-LTx IRI.

Ferroptosis is a regulated form of cell death that results from iron-dependent lipid peroxidation without exhibiting morphological and biochemical characteristics of typical cell death such as cell shrinkage, mitochondrial fragmentation, or nuclear condensation(24). The hallmarks of ferroptosis include dysfunction of lipid peroxide clearance, the presence of redox-active iron as well as oxidation of polyunsaturated fatty acid (PUFA)-containing phospholipids. Depletion of intracellular glutathione and inactivated activity of GPX4 leads to ferroptosis of affected cells, as generation of lipid peroxides cannot be abolished by the GPX4-catalyzed reduction reaction. Moreover, excessive iron accumulation and conversion to Fe^2+^ triggers ferroptosis by mediating the generation of cellular lipid hydroperoxides and lipid peroxidation thereby causing cell death.

Ferroptosis contributes greatly to the pathogenesis of lung diseases, including chronic obstructive pulmonary disease, asthma, cystic fibrosis, pulmonary fibrosis, acute respiratory distress syndrome, and lung cancer (11). Similarly, results from our study shows significantly increased iron levels in lungs that underwent IRI as well as increased levels of oxidized lipids that are a hallmark of ferroptosis. A recent study showed that oxidation of phosphatidylcholines (PC) during IRI generate bioactive phospholipid intermediates that disrupt mitochondrial bioenergetics and calcium transients and provoke wide spread cell death through ferroptosis (25). Similarly, our findings show that the levels of intact PC species were reduced and oxidized PC were increased in lung IRI, establishing ferroptosis in lung IRI. Ferroptosis can result in the accumulation of immune cells and promote the release of proinflammatory cytokines, which can aggravate tissue injury forming a self-amplified loop, which further promotes organ damage (26). Our results demonstrate improved lung functions, reduced inflammation and mitigation of ferroptosis-specific lipid peroxidation following pharmacological inhibition of ferroptosis inhibitor as well as by SPMs such as LxA_4_, indicating an important cross talk between ferroptosis and inflammation in lung IRI.

There is increasing evidence suggesting that lipoxins can potently modulate inflammation within the lung (27). Lipoxins can mitigate the generation of ROS and pro-inflammatory cytokines thereby promoting the resolution of tissue inflammation. In particular, LxA_4_ elicit its actions by binding to formyl peptide receptor-2 (FPR2, ALX/FPR2), a GPCR that transduces downstream signaling to modulate inflammation. FPR2 is expressed in the surface of numerous cell types like neutrophils, monocytes, macrophages, T-cells, enterocytes, airway epithelium, fibroblasts and intestinal epithelial cells (28, 29). Studies have shown that activation of FPR2 receptors offers protection in animal models of lung injury (30, 31). The interaction of LxA_4_ with FPR2 to provide protection was recently shown in a mice model of endotoxin induced acute lung injury (32). Lipoxins promote FPR2 homodimerization, driving a pro-resolving signaling mainly mediated by the inhibition of NF-kB-related pro-inflammatory response, the modulation of intracellular calcium influx and the induction of several beneficial effects (28). Our results show that deletion of FPR2 abolished the protective effects of LxA_4_ in lung IRI confirming that the protective effects of LxA_4_ are FPR2 dependent.

Previously, we have reported that LxA_4_ expression in BAL increases significantly in post-LTx patient samples(4). However, the endogenous levels of bioactive isoforms of SPMs, such as Resolvin D1, Maresin 1 and LxA_4_ may not be sufficient to overcome the initial allograft insult and inflammatory milieu that can lead to significant IRI and tissue damage. Other studies have reported that diminished levels of LxA_4_ are detected in the induced sputum of patients with severe asthma that is characterized by irreversible airway remodeling, including increased collagen deposition and smooth muscle proliferation in the bronchial wall (27). LxA_4_ has been shown to lower transforming growth factor-β (TGF-β) levels in bleomycin-induced pulmonary fibrosis and exhibits an anti-fibrotic effect (33). Recently, the efficacy of LXA_4_ to protects against erastin induced ferroptosis have been studied in spinal cord neurons (10). In post-LTx IRI, the role of LxA_4_-mediated signaling remained to be deciphered and thus we determined if exogenous LxA_4_ protects against ferroptosis and attenuates lung IRI. Although LxA_4_ can mediate tissue protective signaling via other mechanisms such as PPAR-γ or decreasing the production of leukotrienes in other disease models, our results suggest that LxA_4_-mediated attenuation of lung IRI is regulated via activation of NRF2 in ATII cells.

The transcription factor NRF2 regulates the coordinated expression of the antioxidant response element (ARE)-related genes that mediates the expression of phase II antioxidant enzymes, such as heme oxygenase-1 as well as other enzymes related to glutathione synthesis (34). Of particular importance is the fact that the GPX4 synthesis and function, intracellular iron homeostasis, and lipid peroxidation clearing can all be mediated by NRF2 target genes (35). Previous studies have shown that NRF2 stimulation can abrogate response to ferroptosis activators, and NRF2 activation can inhibit ferroptosis to protect against intestinal IR-induced acute lung injury (21, 36). In this study, we found that LxA_4_ mediated protection against pulmonary dysfunctions and ferroptosis were lost in *Nrf2^−/−^* mice. This suggests that protective effects of LxA_4_ in lung IRI is mediated by both FPR2 receptor binding and NRF2 nuclear translocation pathways, which could function independently. Moreover, *Nrf2^−/−^* mice exhibited marked increase in pulmonary dysfunction and inflammation compared to wild type mice after IRI. Taken together, these data suggest that NRF2 deficiency can further aggravate lung IRI.

As the interface between the host and the external environment, the epithelium plays an important role in protecting the lungs from environmental insults and maintaining the homeostasis of the respiratory system. Pulmonary IRI can cause cellular breakdown and death of lung epithelial tissue, which may contribute to the magnitude and duration of pulmonary dysfunction seen after LTx (37). ATII cells are highly metabolically active under normal physiological conditions. Further, ATII cells have higher levels of phospholipids than normal cells making it more susceptible to lipid peroxidation. Based on this, it is speculated that ATII cells exhibit increased susceptibility to ferroptosis compared to other types of lung cells (38). The results from our scRNA seq analysis shows that ferroptosis in ATII cells is involved in chronic rejection in patients diagnosed with CLAD. One of the key aspects of epithelial cell-mediated injury is release of potent chemokine, CXCL1, that is copiously secreted after exposure to iNKT cell-produced IL-17A and alveolar macrophage-dependent TNF-α, during lung IRI as we have previously reported (39). ATII cell-secreted CXCL1 is markedly involved in chemotaxis and transendothelial migration of neutrophils which secrete MPO to cause alveolar damage during lung IRI. The mitigation of ATII cell-dependent CXCL1 and downstream neutrophil infiltration by LxA_4_/FPR2-dependent NRF2 activation and protection against ferroptosis, represents a previously uncharacterized molecular aspect in this study for prevention of post-LTx PGD.

There are a few limitations of this study. The experimental model employing hilar ligation does not fully replicate all aspects of clinical LTx such as organ preservation or the use of immunosuppression. Therefore, we confirmed our findings using the orthotopic LTx model which involves cold preservation and demonstrated the protective aspects of LxA_4_ treatment. Another clinically important caveat refers to the progression of PGD to chronic rejection which was not delineated in the present study. However, the synergy of molecular aspects of ferroptotic signaling between the murine models of PGD, *in vitro* ATII cell-specific signaling, and scRNA seq data in CLAD patients signifies the relevance of excess iron-mediated ATII cell death from initiation of early lung IRI to chronic allograft rejection in post-LTx cohorts. Collectively, these results imply a significant association between alterations in immune metabolomics involving lipidomic and failed resolution due to ferroptosis-related pathway in post-LTx IRI.

In summary, we have demonstrated the involvement of ATII cell-mediated ferroptosis in mediating lung IRI and the therapeutic efficacy of exogenous LxA_4_ administration in mitigating PGD. Future investigations should focus on the use of combined strategy of bioactive isoforms of SPMs such as Resolvin D1 and Maresin-1 along with LxA_4_ as a multi-faceted approach to target immune metabolism for mitigation of post-LTx PGD to chronic rejection for a viable clinical translation.

## Author contributions

AKS designed the study. DJMK, VL, ZT, MJWS, MK and AKS performed experiments. DJMK, VL, MJWS, CA, GRU and AKS analyzed results. GC performed single cell sequencing analysis. MK and TJG performed lipidomic analysis. DJMK and AKS prepared the manuscript with input from all authors.

## Acknowledgments

This work was supported by David and Kim Raab Foundation (AKS) and National Institute of Health (NIH) RO1 HL140470-0181 (CA). The authors have no conflicts of interest to disclose.

