## Supplementary Materials and Figures for "Lipoxin A_4_/FPR2 signaling mitigates ferroptosis of alveolar epithelial cells via NRF2-dependent pathway during lung ischemia-reperfusion injury"

**Table of Contents:**

1. Materials and Methods
2. Supplementary Figure(s) and Table(s)
    a. Supplementary Figure S1
    b. Supplementary Figure S2
    c. Supplementary Figure S3
    d. Table S1
    e. Table S2
    f. Table S3

### Materials and Methods

#### *Human LTx single-cell RNA sequencing analysis*

We conducted analysis on single cell RNA sequencing (scRNA-seq) data of lungs explanted from four patients with chronic lung allograft dysfunction (CLAD) and three normal donors (DT). The original study has explained data of 40 cell populations in epithelial, endothelial, lymphoid, myeloid and mesenchymal lineages (GSE224210) (1). In this study, we focused specifically on alveolar epithelial type 1 and type 2 cell populations. These populations were filtered based on their original annotation, and we further analyzed cell clustering with the following steps: Data was analyzed using Seurat v5 after applying 'nFeature_RNA > 250 & nFeature_RNA < 4000 & percent.mt < 15' for quality control, 14,371 cells were included in further analysis (2). Using the 'LogNormalize' method data was normalized and CLAD and DT data were integrated using 30 identified anchors. Based on the top 20 principal components (PCs), the shared nearest neighbor (SNN) graph and Louvain algorithm were applied to identify 18 cell clusters, with a resolution parameter of 0.5. Using Uniform Manifold Approximation and Projection (UMAP) embeddings, these cell clusters were visualized in a dimension-reduced space. Cell clusters were annotated using Azimuth (https://satijalab.org/azimuth) based on The Human Lung Cell Atlas scRNA-seq data reference. Further, the CLAD data were compared to control data employing Wilcoxon Rank Sum test and differentially expressed genes were identified by Log2FC>0.25, min.pct>0.01 and Bonferroni corrected p-value<0.05.

Sequencing analysis identified 5,584 genes in ATII epithelial cells, of which 1,839 were differentially expressed genes (DEGs) in CLAD vs. DT (Table S1). We cross-referenced these genes with a curated list of ferroptosis-related genes (FRGs) generated from a GeneCards® and PubMed® query (ferroptosis; Table S2). Using this approach, we ensured precise identification of ferroptosis related genes within the dataset.

#### *Lung IRI model*

An *in vivo* hilar ligation model was used for lung IRI in 8-12 week old male C57BL/6 (wild-type; WT) mice (Jackson laboratory, Bar Harbor, ME), *Nrf2*$^{-/-}$ mice (Jackson laboratory, Bar Harbor, ME) and *Fpr2*$^{-/-}$ mice (Idorsia Pharmaceuticals, Switzerland), as previously described (3). Mice were provided drinking water and standard chow diet *ad libitum*. All procedures were conducted in accordance to the Institutional Animal Care and Use Committee of the University of Florida (protocol # 202110465). In brief, mice were anesthetized with inhaled Pivetal® Isoflurane (Patterson Veterinary, Gainesville, FL) intubated with PE-60 tubing and connected to a pressure-controlled ventilator (Harvard Apparatus Co, South Natick, MA). Mechanical ventilation with room air was done at 150 strokes/min, 0.5 cc stroke volume, and peak inspiratory pressure <20 cm $H_2O$. The mice underwent 1hr of ischemia followed by 6hr of reperfusion while sham animals received the same surgery but without hilar occlusion. Mice were treated intratracheally with normal saline (NS) or $LxA_4$ (100 ng/kg; Cayman Chemicals, Ann Arbor, MI) 1 h prior to ischemia. The selection of $LxA_4$ dosage was based on previous studies from our group and published literature (3, 4). To determine the effect of pharmacological inhibition of ferroptosis on lung IRI, animals were intraperitoneally (i.p.) injected with liproxstatin-1 (50 mg/kg; Cayman Chemicals) or vehicle control (DMSO) 1hr before ischemia (5, 6). To minimize potential ventilator-induced injury, mice were extubated and returned to their cage during both the ischemic and reperfusion periods.

#### *Lipidomic analysis*

Lung samples were extracted using the Bligh-Dyer extraction method (7). Subsequently, liquid chromatography was performed on an ultra-high-performance liquid chromatography (UHPLC) system (Vanquish;Thermo Scientific, San Jose, CA) and samples were analyzed using ACQUITY UPLC BEH C18 1.7 µm, 2.1 mm X 50 mm column with ACQUITY UPLC BEH C18 VanGuard Pre-column 1.7 µm, 2.1 mm X 5 mm. Also, a gradient program consisting of mobile phase A (60:40 acetonitrile/water) and mobile phase B (90:10 isopropanol/acetonitrile), each containing

10 mmol/L ammonium formate and 0.1% (vol/vol) formic acid, was used for sample analysis. The ultra-high-performance liquid chromatography (UHPLC) system was coupled to a mass spectrometer (Q-Exactive Orbitrap; Thermo Scientific) for chromatographic separation and mass spectral measurement of lipids in positive and negative ion mode, using full-scan and iterative exclusion tandem mass spectrometry. The software LipidMatch Flow was deployed for peak picking, blank filtering, annotation, and combining positive and negative ion polarity. Additionally, LipidMatch Normalizer was used for semi quantitation (8).

***Pulmonary function analysis***

Pulmonary function was assessed using a buffer-perfused, isolated mouse lung system (Hugo Sachs Elektronik, March-Huggstetten, Germany) as previously described (9). Briefly, mice were anesthetized with ketamine and xylazine after the reperfusion period, and a tracheostomy was performed. Using a MINIVENT mouse ventilator (Hugo Sachs Elektronik, March-Huggstetten, Germany), animals were ventilated with room air at 100 breaths/min at a tidal volume of 7μl/g body weight with a positive end expiratory pressure of 2 cm $H_2O$. The lungs were perfused with Krebs-Henseleit buffer (Sigma-Aldrich, St. Louis, MO) containing 2% albumin, 0.1% glucose and 0.3% HEPES (335–340 mOsmol/kg $H_2O$) at a constant flow of 60μl/g body wt/min. For equilibration, the lungs were maintained on the system for a 5-minute period before recording data for an additional 5 minutes.

***Murine BAL analysis***

After finishing pulmonary function measurements, left lungs were lavaged with 0.3 mL PBS. The BAL fluid was normalized for return volume collected, then centrifuged at 4°C (500*g* for 5 min). The supernatant was collected and stored at –80°C until further analysis.

***Prussian blue staining***

Perls’ Prussian blue staining was performed in lung tissue using the Iron Stain Kit (HT20-1KT, Sigma-Aldrich, St. Louis, MO, USA), according to manufacturer’s instructions. Paraffin-embedded tissue samples were rehydrated through a series of graded alcohol and $H_2O$, stained with Perls’ staining solution for 20 min, rinsed with deionized water, counterstained with nuclear fast red, rinsed with deionized water and then rapidly dehydrated. Images were taken with 40X magnification using a Nikon microscope equipped with a digital camera and Elements BR software.

***Estimation of iron content***

Iron concentration in lung tissue extracts was measured using an iron assay kit, per the manufacturer's instructions (Millipore Sigma, St. Louis, MO).

***Cytokine and chemokine protein analysis***

Cytokine and chemokine protein content in BAL fluid and cell culture supernatants were measured using a mouse-specific Milliplex Cytokine Assay panel (Millipore Sigma, Burlington, MA).

***Myeloperoxidase (MPO) measurement***

Myeloperoxidase (MPO) in murine BAL fluid was measured to assess neutrophil infiltration into the alveolar space using an MPO ELISA kit (Cell Sciences, Canton, MA) as instructed by the manufacturer.

***Immunohistochemistry***

Immunostaining of the murine lungs was performed to identify neutrophils. Left lungs were harvested following 6hr reperfusion, fixed overnight in 10% neutral buffered formalin (Sigma-Aldrich, St. Louis, MO), embedded in paraffin and sectioned at 5 μM. Immunostaining was performed to identify neutrophils with purified rat anti-mouse Ly-6G (BD Biosciences, San Jose, CA). Alkaline phosphatase-conjugated anti-rat immunoglobulin G (Vector Laboratories, Burlingame, CA) was used as secondary antibody, and the signals were detected with Vector®

Red substrate kit (Vector Laboratories). Mayer's Hematoxylin (Thermo Fisher Scientific, Waltham, MA) was used to counterstain the sections. Neutrophils for each lung section were counted by a blinded reviewer in five random fields. The counts were averaged per tissue sample and included all cells in peripheral lung tissue (3). Iron from lung sections were detected using iron stain kit (Millipore Sigma, Burlington, MA) as per the manufacturer's instructions.

***Deceased Cardiac Donor (DCD) and Orthotopic Lung Transplantation***

Orthotopic lung transplants were carried out on DCD donors using Balb/c donor and C57Bl/6 recipient mice, as previously descried (9). In brief, Balb/c donor mice were anesthetized by injection of ketamine and xylazine intraperitoneally (0.01 mg/g) and ventilated with isoflurane and oxygen mixture. After median laparosternotomy, heparinization was done by intravenous injection of 100 U/body, and cardiac arrest was induced by intravenous injection of potassium chloride at 4 mg/body. Following the induction of cardiac arrest, 100 μl NS or $LxA_4$ (100 ng/kg) were intratracheally administrated and the ventilated donor mice were monitored for a 60 min period of "no-touch" warm ischemia. After warm ischemia, donor lungs were flushed through the main pulmonary artery with 2 mL of 4°C Perfadex®, harvested and stored for 18hrs in Perfadex® at 4°C. Donor left lungs were prepared for transplantation and just prior left orthotopic vascularized LTx, either 30 μL NS or $LxA_4$ was given via left bronchial. Utilizing cuff techniques, left lungs were transplanted into C57Bl/6 recipient mice.

***Arterial blood gas measurements***

At the end of 24hrs reperfusion period after LTx, mice were placed on mechanical ventilation (10 μL/g tidal volume, respiratory rate 120/min, PEEP 3 cm $H_2O$) with 100% oxygen for 5 min. Blood samples were collected from the left ventricle into heparinized syringes to measure arterial blood gas. A VetScan i-STAT 1 handheld analyzer and CG4+ cartridges were used for measuring arterial blood gases (Abaxis, Union City, CA) (3).

***In vitro hypoxia-reoxygenation (HR)***

Alveolar type II epithelial (MLE12; ATCC) cells were cultured in DMEM F-12 medium (Gibco), containing 1% penicillin/streptomycin (Invitrogen, Carlsbad, CA) and 10% FBS in 6-well culture plates at 37 °C and 5% $CO_2$. Culture plates were exposed to hypoxia/reoxygenation (HR) by placing them in a humidified, sealed hypoxic chamber (Billups-Rothenberg, Del Mar, CA) and purged with 95% $N_2$ and 5% $CO_2$ for 25 min as described previously (3). The sealed chamber was then placed in a cell culture incubator at 37 °C for 3hrs hypoxia, after which the chamber was opened and reoxygenation was achieved by transferring the plates from the hypoxic chamber into a normoxic humidified incubator (37°C, 5% $CO_2$) for an additional 3hrs. FPR2 receptor knockdown was achieved in MLE12 cells via transfection of FPR2 siRNA (1μM; Dharmacon) using Lipofectamine RNAiMax transfection reagent (Thermo Fisher Scientific), as previously described (10). $LxA_4$ (300 nM; Cayman Chemicals) was added to cell culture media 1hr prior to initiation of hypoxia. Selected dose of $LxA_4$ was based on previous studies. Cell lysates and supernatant media were harvested for measuring ferroptosis markers. Quantification of DCF dye in ATII cells was performed using an OxiSelect Intracellular ROS assay kit as instructed (Cell Biolabs, San Diego, CA). All DCF solutions were protected from light to prevent light-induced auto-oxidation. Lipid peroxidation (MDA; Millipore Sigma, St. Louis, MO) and glutathione (GSH; Cayman Chemicals, Ann Arbor, MI) were measured in cell culture or tissue extracts using colorimetric assay kits, per the manufacturer's instructions. Separate ATII cell cultures were also treated with erastin (10 μM; ferroptosis activator; Cayman Chemicals, Minneapolis, MN) with/without LxA4 (300nM; Cayman Chemicals) and analyzed for MDA and GSH expressions in ATII culture extracts after 6hrs. NRF2 transcription factor activation in nuclear extracts was measured using a colorimetric assay kit (Abcam, Cambridge, UK).

***Statistical analysis***

Values are presented as the mean ± standard error of the mean (SEM) and statistical evaluation was performed with GraphPad Prism 10 software (GraphPad, La Jolla, CA). Parametric one-way

ANOVA followed by Tukey's multiple comparison test was performed to compare differences between three or more groups, and Mann-Whitney test was used for pair-wise comparisons of groups. A value of $P < 0.05$ was considered statistically significant.

**Figure S1**

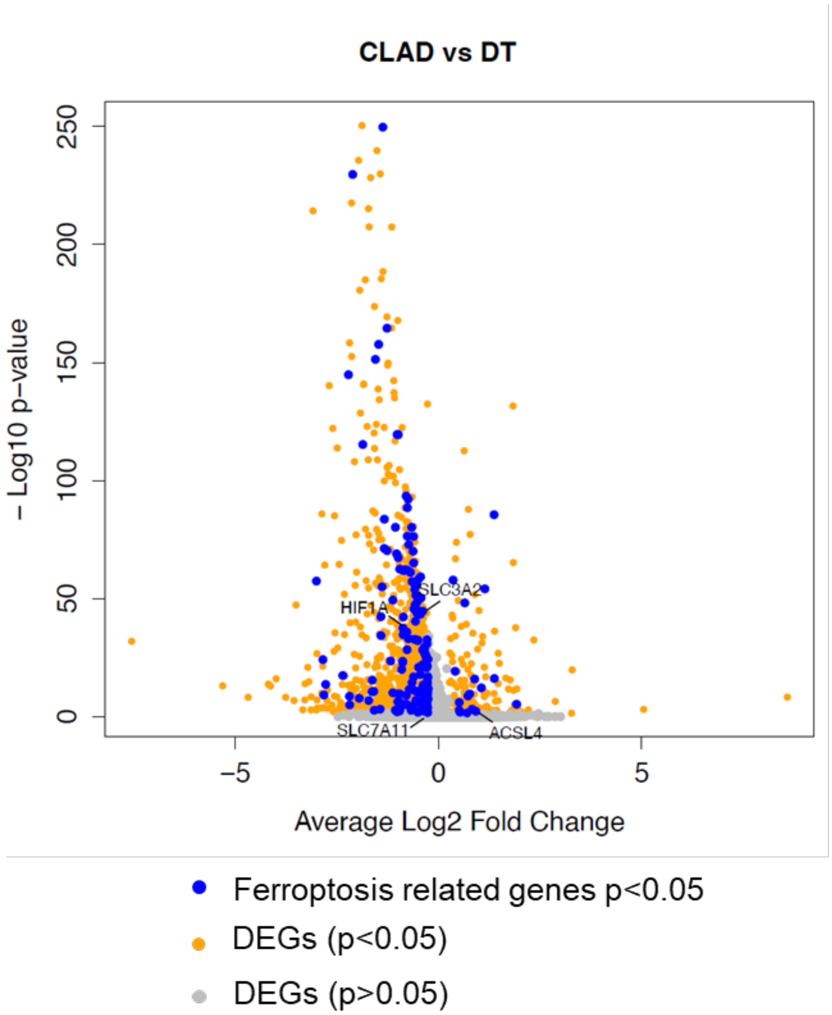


**Figure S1**: Volcano plot illustrating the differentially expressed genes (DEGs) within ATI cells. Ferroptosis related genes with a significant differential expression ($p<0.05$) are depicted as blue dots while other significant DEGs ($p<0.05$) are represented by orange dots. Genes lacking statistical significance ($p>0.05$) are represented by grey dots.

**Figure S2**

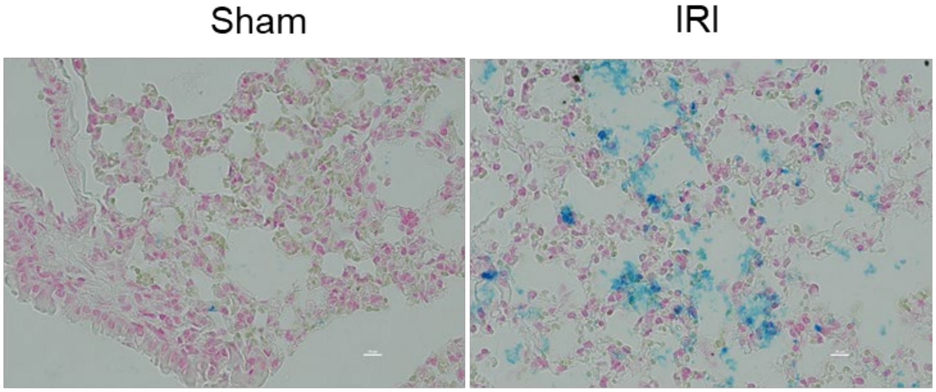


**Figure S2**. Representative images of Prussian blue staining from mice lung sections after sham or ischemia reperfusion injury (IRI). Scale bars indicate 10 µm.

**Figure S3**

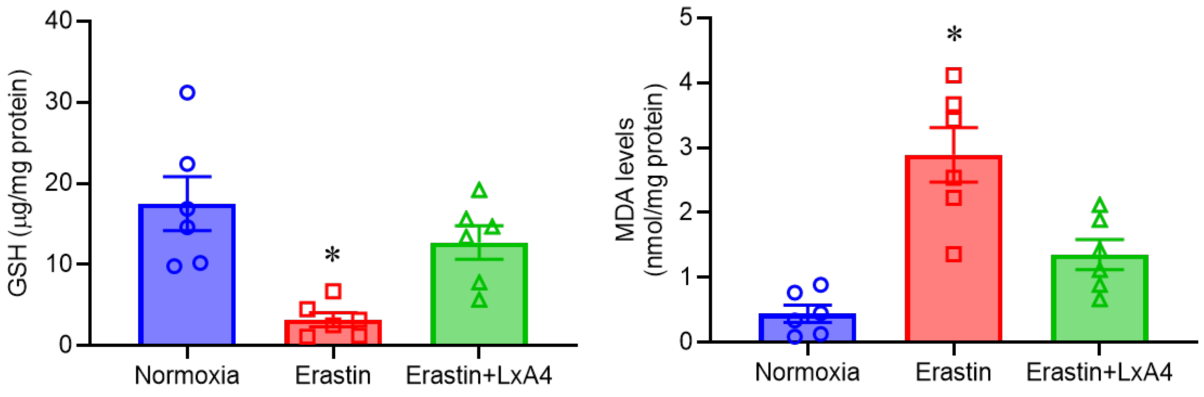


**Figure S3**. A significant decrease in GSH and increase in MDA was observed in ATII cells after erastin treatment which was abolished by LxA4 treatment. *p<0.02 vs. other groups.
